# In a model of parasite-mediated exhaustion, stem-like CD8 T cells differentiate into an unconventional intermediate effector memory subset

**DOI:** 10.1101/2024.10.30.621158

**Authors:** Magali M. Moretto, Keer Chen, Christina Cox, Jie Chen, Imtiaz A Khan

**Affiliations:** Department of Microbiology, Immunology, and Tropical Medicine, The George Washington University, Washington DC, USA

**Keywords:** Toxoplasma gondii, CD8 T cells, memory T cells, stem-like T cells, intermediate CD8 T cells, exhaustion

## Abstract

CD8 T cell exhaustion has been reported in mice susceptible to *Toxoplasma gondii* infection. While the differentiation of CD8 exhausted subsets has been extensively reported, most of these studies have been conducted in chronic viral and cancer models. During chronic *T. gondii* infection, phenotypic and transcriptomic analyses of the polyclonal antigen-specific CD8 T cell response characterize four populations based on KLRG1 and CD62L expression. Pop1 (KLRG1^+^CD62L^lo^) bears the attributes of a terminal effector subset, and pop2 (KLRG1^-^CD62L^lo^) is similar to effector memory CD8 T cells. Akin to chronic viral infection and cancer systems, pop3 (KLRG1^-^CD62L^hi^) exhibits the characteristics of stem-like progenitor CD8 T cells (high Tcf7, Slamf6, and Cxcr5 expression), whereas pop4 (KLRG1^+^CD62L^hi^) closely resembles a transitory subset (elevated Tbx21, low Tcf1, and Tox expression). During chronic viral infection, the stem-like progenitor CD8 T cells transition into a terminally differentiated exhausted subset via an intermediate population. However, in our system, pop3 generates pop4, which does not convert into a conventional terminally differentiated exhausted subset but instead transitions into effector pop1. Notably, during the chronic phase of the infection, pop1 cannot retain its functionality, irrespective of its origin, which may hamper its ability to control reactivation. Our observations emphasize that the differentiation of exhausted CD8 T cells in non-viral infections, like chronic toxoplasmosis, follows a different pattern than established models and highlights the need to develop new immune strategies better tailored for a broad range of pathogens.

## Introduction

CD8 T cell immunity plays a critical role in protection against intracellular pathogens and malignancies (1, 2). Upon TCR engagement and co-stimulation by antigen-presenting cells, naïve CD8 T cells differentiate into effector and memory precursor effector (MPEC) CD8 T cells (3). Terminally differentiated effector CD8 T cells provide immediate protection and contract once pathogens are controlled (4). On the other hand, memory CD8 T cells derived from MPEC are long-lived and provide long-term protection against recurring infections (5). Memory CD8 T cells have been categorized into central memory (T_CM_) and effector memory (T_EM_) populations based on CCR7 and CD62L expression (6). T_CM_ (CD62L^hi^CCR7^hi^) exhibits increased longevity, greater homeostatic proliferation, and recall response (7). In comparison, T_EM_ (CD62L^lo^CCR7^lo^) has a shorter life span and displays increased cytokine production and cytotoxic activity (7). This paradigm became more complex as additional populations like tissue-resident memory, stem-like memory, and intermediate effector memory CD8 T cells have been identified (8, 9). In the context of chronic infections or cancer, constant antigenic stimulation impairs the memory CD8 response, as a result, cells become exhausted and subsequently undergo apoptosis (10). Interestingly, in these conditions, stem-like CD8 T cells, which are long-lived, due to their plasticity, can replenish the CD8 T cell populations. These cells are prime immunotherapeutic targets and critical for controlling chronic infection and cancer (11).

Toxoplasmosis, caused by the obligate intracellular parasite *T. gondii*, is a chronic infection that affects up to a third of the world’s population (12). The parasite induces a strong inflammation and both innate and adaptive immune responses are critical for controlling the acute infection. Long-term protection is predominantly dependent on the CD8 T cell memory population, and their polyfunctional response is essential for preventing reactivation (13). While BALB/c mice (H-2L^d^) are resistant to toxoplasmosis, C57BL6 (H-2K^b^) are susceptible and develop encephalitis (14). In a susceptible model, the inability to control the chronic infection is attributed to a loss of CD8 T cell function caused by exhaustion (15) similar to chronic viral infection and cancer (11, 16). In these systems, memory CD8 T cells are comprised of a stem-like progenitor exhausted, an intermediate exhausted but functional, and a terminally differentiated exhausted dysfunctional CD8 subset (17). Notably, in chronic non-viral infections, stem-like memory, and intermediate effector memory CD8 T cell subsets are not well studied (18, 19).

Using susceptible C57BL6 mice, the present study demonstrates that the memory CD8 T cell response after *T. gondii* infection cannot be categorized based on CD62L and CCR7 expression. However, using a clustering algorithm, CD8 T cells can be divided into four subsets with differential KLRG1 and CD62L expression: KLRG1^+^CD62L^lo^ (pop1), KLRG1^-^CD62L^lo^ (pop2), KLRG1^-^CD62L^hi^ (pop3), and KLRG1^+^CD62L^hi^ (pop4). Pop1 resembles a classic effector subset, pop2 is similar to T_EM_, pop3 appears to be a stem-like memory subset, and pop4 resembles an intermediate effector/memory population. Unlike chronic viral and cancer systems, there is no linear differentiation from a stem-like CD8 T cell subset to an intermediate population that can subsequently convert into a terminally differentiated exhausted phenotype during *T. gondii* infection. Instead, stem-like pop3 can directly generate pop2 and pop4 with different functional, phenotypic, and transcriptional attributes. Our results demonstrate that in a susceptible encephalitis model of *T. gondii* infection, the differentiation of CD8 T cell populations is likely mediated by pathogen-specific cues, highlighting the need for broader immunotherapeutic strategies.

## Results

### CD62L and KLRG1 expression defines four antigen-experienced CD8 T cell subsets in *T. gondii*-infected animals

Antigen-experienced (Ag^+^) memory CD8 subsets were analyzed using a published surrogate marker strategy (CD44^hi^CD11a^hi^)(20, 21) since there is currently only one tetramer (tet^+^: Tgd057) that recognizes a small fraction of Ag^+^ cells in the spleen of chronically infected C57BL6/J mice (22) (Fig. S1A). The surrogate marker approach facilitates the identification of polyclonal CD8 T cell response in *T. gondii* infected animals while detecting only 10% or fewer CD8 T cells from naïve mice (Fig. 1D). Furthermore, parallel analysis of Ag^+^ and tet^+^ (Tgd057^+^) CD8 T cells at 6 weeks post-infection (p.i.) demonstrates that the majority of tet^+^ cells in the brain, liver, and spleen are predominantly CD44^hi^CD11a^hi^ (Ag^+^)(Fig. S1A). These findings confirm previous observations reporting that tet^+^ CD8 T cells comprise a higher percentage of IFNγ^+^ cells than CD11^lo^CD44^lo^ CD8 T cells (21). Our data also substantiates a previous study (23) showing that *T. gondii* infection does not generate classical T_EM_ (CD62L^lo^CCR7^lo^) and T_CM_ (CD62L^hi^CCR7^hi^) memory subsets observed in other models (6). As shown in figure S1B-C, a significant percentage of memory (CD127^hi^) Ag^+^ CD8 T cells from the brain, liver, and spleen of infected mice labeled with CCR7 and CD62L fall outside the established T_CM_/T_EM_ model. Even though KLRG1, a marker commonly associated with effector CD8 T cells (24), can be expressed by some memory subsets (25, 26), an unexpectedly high frequency of CD62L^lo^CCR7^lo^ and CD62L^hi^CCR7^hi^ CD8 T cells from *T. gondii* infected mice express this marker (Fig. S1D). Therefore, unbiased clustering with FlowSOM algorithm was applied to categorize Ag^+^ CD8 T cells based on four well-described CD8 T cell subset markers: CD62L, KLRG1, CD127, and CCR7. As shown in figure 1A, the Ag^+^ CD8 T cells were clustered into four populations with differential expression patterns of CD62L and KLRG1 (Fig. 1B-C): KLRG1^+^CD62L^lo^ (pop1), KLRG1^-^ CD62L^lo^ (pop2), KLRG1^-^CD62L^hi^ (pop3), and KLRG1^+^CD62L^hi^ (pop4) (Fig. 1D). Interestingly, these subsets have been previously described in the peritoneal exudate, spleen, and brain from mice infected with a vaccine strain of *T. gondii* (22). This report evaluated the role of IL-12 in the regulation of memory CD8 T cells, but the subsets were not further characterized. Therefore, in this study, expression of markers associated with memory and effector responses (CD127, CCR7, CD62L, KLRG1, CX3CR1) by Ag^+^ CD8 T cells at week 5 p.i. was visualized by flow cytometry using the dimensionality reduction algorithm TriMap (Fig. 1E). When overlayed on the TriMap plot, pop1-4 effectively comprise most of the Ag^+^ CD8 T cell population with minimal overlap (Fig. 1F). At week 5 p.i., KLRG1^+^ pop1 represents the largest subset (in percentage and frequency) while the other three subsets are equally represented (Fig. 1G). All four subsets were detected during the acute as well as chronic phase of infection (Fig. 1H). As expected, pop1 (KLRG1^+^CD62L^lo^) peaks at week 2 p.i., when the parasite is actively replicating and decreases slowly during the chronic phase. However, this subset remains the largest population detected at weeks 5 and 8 p.i. (Fig. 1H). This could be attributed to the unexpected replication of the parasite during the chronic phase of the infection (27). Likewise, the number of cells in pop2 (KLRG1^-^CD62L^lo^) decreases over the course of the infection (Fig. 1H), while pop3 (KLRG1^-^CD62L^hi^) and pop4 (KLRG1^+^CD62L^hi^) increase as the infection progresses to chronicity. As observed in the spleen, pop1 is predominant in the brain and liver at week 2 p.i., when the parasites are disseminating to the different tissues, as well as during the late chronic phase of the infection (Fig. S2). Unlike splenic tissue, the brain and liver display elevated frequency of pop1 and pop2 while pop3 and pop4 are less represented as previously published (22).

**Figure 1.**
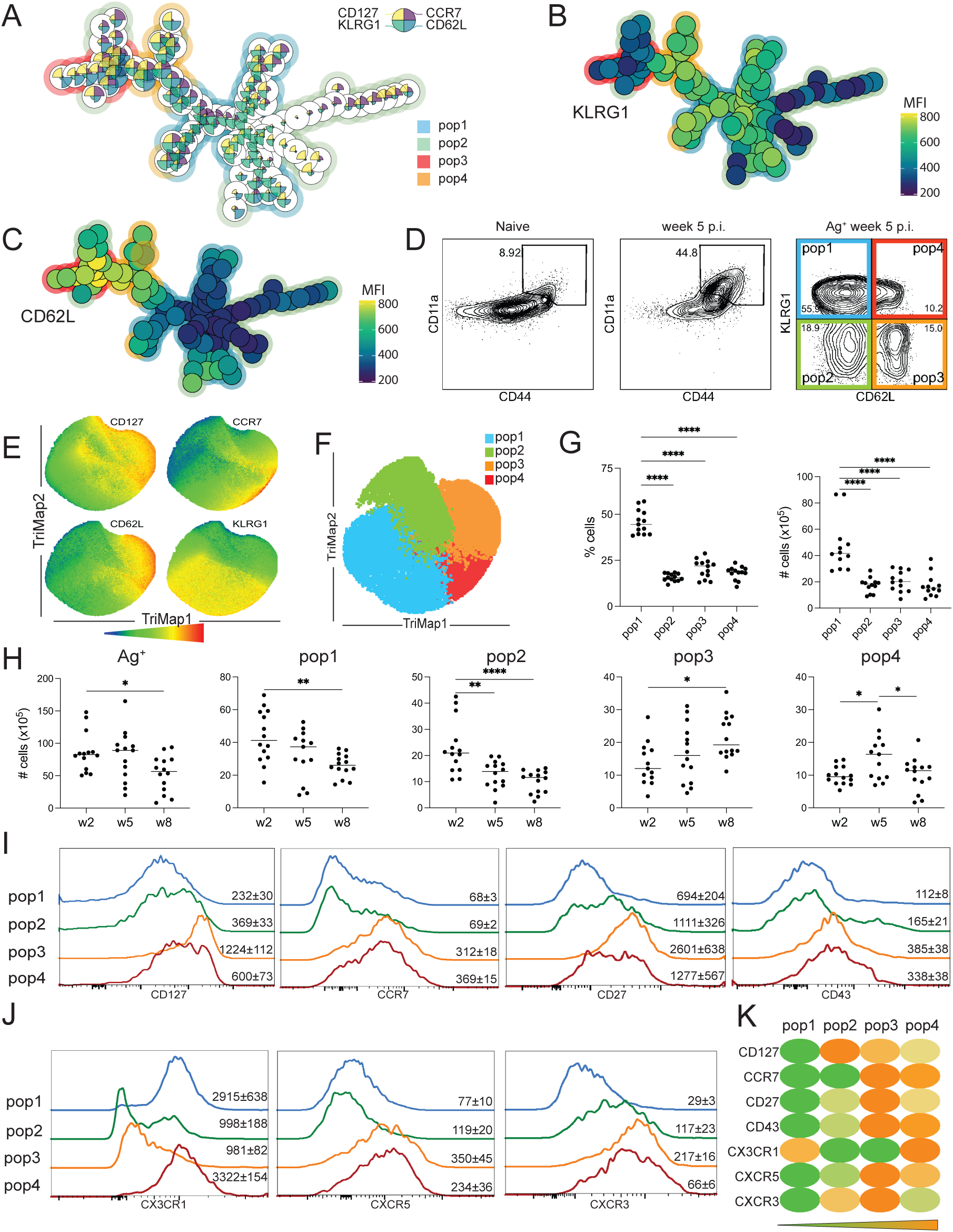
Phenotypic characterization of CD8 T cell subsets after *T. gondii* infection. C57BL6/J mice were infected with 10 cysts of *T. gondii* (strain Me49) via oral route. A) FlowSOM minimal spanning tree of Ag^+^ (CD44^hi^CD11a^hi^) CD8 T cells from the spleen using antibodies against CD127, CD62L, CCR7, and KLRG1 at week 5 p.i. B-C) KLRG1 (B) and CD62L (C) expression overlay on the clustering tree. D) Four populations characterized by KLRG1 and CD62L expression (gated on Ag^+^ CD8 T cells) at week 5 p.i. (pop1: KLRG1^+^CD62L^lo^, pop2: KLRG1^-^CD62L^lo^, pop3: KLRG1^-^CD62L^hi^, pop4: KLRG1^+^CD62L^hi^). E) TriMap representation of CD127, CCR7, CD62L, and KLRG1 expression by Ag^+^ CD8 T cells at week 5 p.i. F) Pop1-4 overlay on the TriMap representation at week 5 p.i. G) Frequency and number of cells for pop1-4 in the spleen at week 5 p.i. H) Kinetic of Ag^+^ and pop1-4 cell numbers at week 2, 5, and 8 p.i. I-J) CD127, CCR7, CD27, CD43 (I) and CX3CR1, CXCR5, and CXCR3 (J) expression by pop1-4 at week 5 p.i. Numbers on histograms represent the mean of the geometric mean fluorescence intensity (gMFI) for one experiment (4 mice) ± SEM for each marker. K) Heatmap of gMFI for markers from (I) and (J) for pop1-4 at week 5 p.i. A-F: Data is representative of at least two experiments with 4 mice per group. G-H: Data presented as the mean of individual mice ± SEM compiled from at least three experiments. *p<0.05, **p<0.01, and ****p<0.001 (one-way ANOVA with Bonferroni post-test).

The different populations in the spleen were further characterized with various memory/activation markers: CD27 promotes long-term survival of CD8 T cells (28), activation associated isoform of CD43 (6), CX3CR1 associated with cytotoxic effector function in memory CD8 T cell subsets (29), CXCR3 regulates stem-like memory and proliferative CD8 T cells, (30), and CXCR5 an effector memory marker in LCMV infection (Fig. 1I-K) (31). Pop1 resembles effector T cells with low expression of CD127, while pop2 displays high levels of CD127 and CXCR3 (Fig. 1I-J). CD127^+^pop3 and pop4, which can be classified as memory subsets, also exhibit CCR7, CXCR5, and CD43. Furthermore, KLRG1^+^ pop1 and pop4 co-express the cytotoxic effector function marker CX3CR1 (Fig. 1J). As shown in figure 1K, pop3 and pop4 display a similar expression pattern that is notably different from pop1 and pop2.

Next, the transcriptional profile of each population was characterized by bulk RNA sequencing during the late chronic phase of the infection when the CD8 T cell response is compromised (week 8 p.i.)(15). The principal component analysis demonstrates that each population displays a unique gene expression signature that differs from the CD8 T cells isolated from naive mice (Fig. 2A). The PCA also reveals that pop1 and pop4 are more alike despite pop4 exhibiting a strong phenotypic resemblance with pop3 (Fig. 1K). Higher coefficient Spearman’s factors are detected within duplicates as well as between pop1 and pop4, underscoring the PCA results (Fig. 2B). Conversely, a lower coefficient factor is observed when comparing pop3 to pop1 or pop4, emphasizing the differences between pop3 and these populations. The Venn diagram confirms that pop4 has more differentially expressed genes (DEGs) in common with pop1 than pop3 when compared to CD8 T cells from naïve mice (Fig. 2C). Next, the top 3800 DEGs between the four populations were divided into six clusters using K-means clustering (Fig. 2C-E, S1 Table). Pop1 and pop4 share a large part of their transcriptome with only 43 DEGs dysregulated (Fig. 2D), including Sell, which encodes CD62L (adjusted p value= 1.07x10^-81^)(Fig. 2E). DEGs that are exclusively upregulated in pop2 belong to cluster 1, which contains markers associated with exhaustion, like Lag-3, Pdcd1, and Cd160 (Fig. 2F). Genes associated with stem-like cells (Tcf7, Cd69, Cxcr5, and Nt5e) (cluster II) are upregulated exclusively in pop3, while those linked to effector T cell differentiation (cluster III) are down-regulated. Cluster IV, with higher expression in pop2 and pop3, is comprised of genes associated with both T cell activation and exhaustion (Cd200, Tigit, Myc, Icos, and Cd27). Genes related to the effector phenotype (perforin, granzymes, T-bet, annexins, and caspases) are predominantly expressed in pop1 and 4 (cluster V), demonstrating that transcriptomic similarities between pop1 and pop4 are strongly linked to a terminal differentiation signature commonly associated with an effector response. Cluster VI is comprised of the 43DEGs that are different between pop1 and pop4. Cluster VIa includes genes with increased expression in pop1 (Lgals3, Plxna1, and Nrp1), while cluster VIb contains those increased in pop4 (Il23r, Gzmm, and Fcgrt). Genes from clusters I-V were then analyzed with Metascape to determine the overrepresented biological pathways. Clusters I, III, IV, and V share several common immune processes (regulation of lymphocyte activation, regulation of cell-cell adhesion, and regulation of lymphocyte proliferation). However, pathways for nucleotide metabolic process and translation at synapse are only associated with cluster II (pop3). Also, this cluster displays elevated genes encoding ribosomal protein, which were recently linked to stem-like CD8 T cells (32). Interestingly, negative regulation of immune response pathway is significantly represented in clusters I, II, and IV, which encompass all four populations.

**Figure 2.**
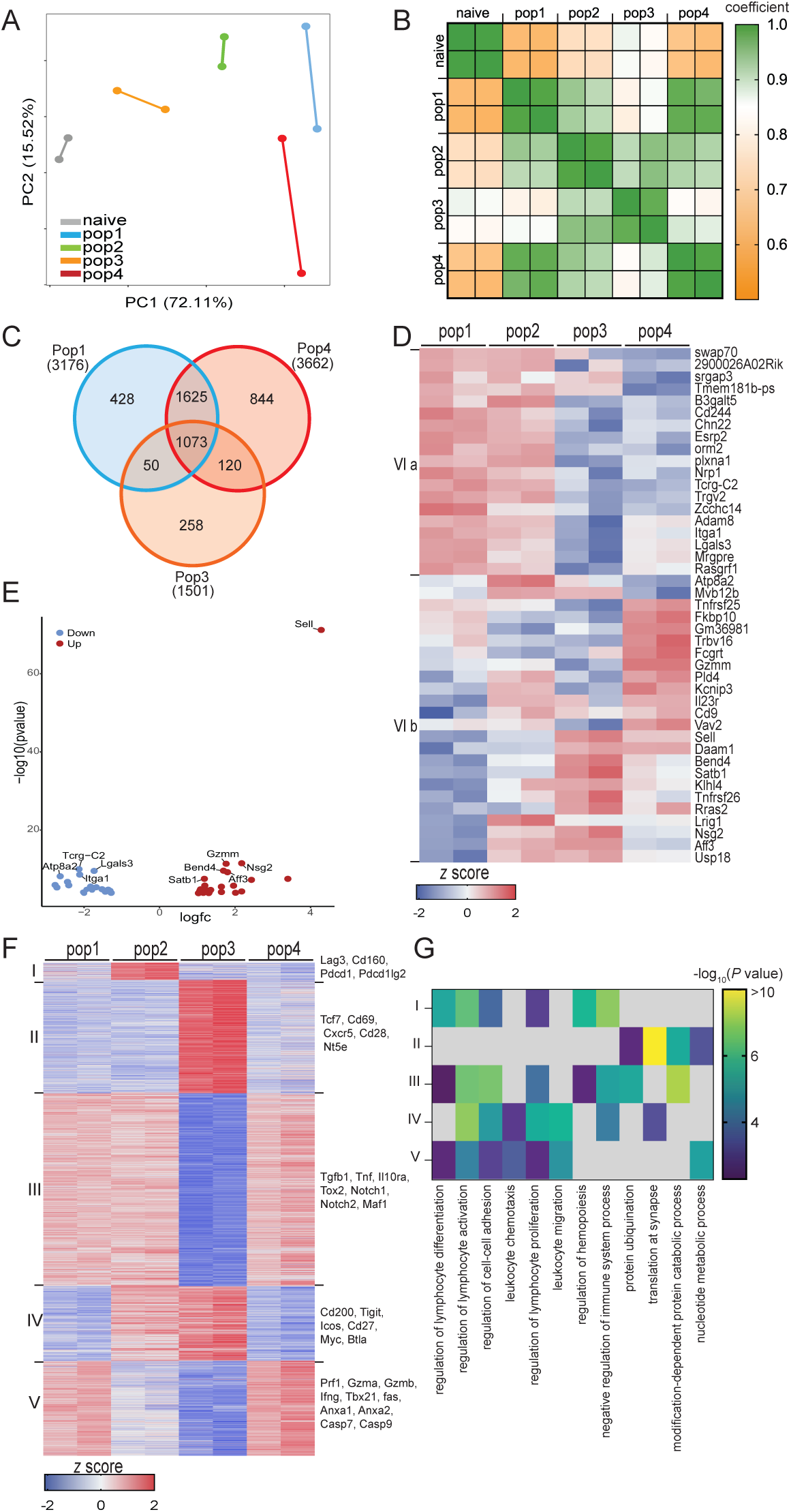
Transcriptomic analysis of CD8 T cell subsets defines four individual populations. C57BL6/J mice were infected with 10 cysts of *T. gondii* (strain Me49) via oral route. At week 8 p.i., CD8 T cell subsets were sorted from the spleen, and RNAseq was performed. A) Principal Component Analysis for pop1-4 and CD8 T cells from naïve mice. B) Heatmap illustrating Spearman’s correlation among individual samples. C) Venn diagram for DEGs from pop1, pop3, and pop4. D) Heatmap showing the relative expression of the 43 DEGs dysregulated between pop1 and pop4 at w8. E) Volcano plot of the 43 DEGs in pop4 compared to pop1. F) Heatmap showing the relative expression of 3800 DEGs between the four populations at w8. G) Metascape analysis indicating the biological processes associated with genes from the clusters defined in (F). D, F: Genes were divided into clusters by K-mean clustering based on expression. Data are from two independent experiments, with samples isolated from 5-7 pooled mice.

Overall, based on phenotypic and transcriptomic analyses, the CD8 T cell response during *T. gondii* infection can be segregated into four different subsets. Pop1 exhibits the hallmark signature of terminally differentiated effector CD8 T cells, and the other three populations display higher CD127 expression associated with memory subsets. pop2 closely resembles T_EM_ (CCR7^lo^CD62L^lo^), and pop3 is similar to stem-like memory CD8 T cells. Although pop4 and pop3 appear phenotypically alike, RNA seq analysis demonstrates that pop4 resembles pop1 with only a 43 DEGs difference.

### Pop3 bears the characteristics of stem-like memory CD8 T cells and gives rise to multiple CD8 T subsets

Next, to determine the plasticity of the different CD8 subsets, adoptive transfer studies were performed. The four populations were isolated from CD45.1 mice at week 5 p.i. and individually transferred to equally infected (5 weeks p.i.) congenic (CD45.2) mice. The recipients were sacrificed 6 days post-transfer and the donor population (CD45.1) was characterized (Fig. 3A). As shown in figure 3 B, a significantly higher number of CD45.1 CD8 T cells is recovered from the mice that received pop3 as compared to the other subsets. The majority of CD45.1 cells recovered from mice transferred with pop1 or pop3 retained their original phenotype, while CD45.1 cells from pop2 recipients partially converted into pop1 (Fig. 3C). Interestingly, the majority of CD45.1 CD8 T cells from pop4 recipients switched to effector pop1.

**Figure 3.**
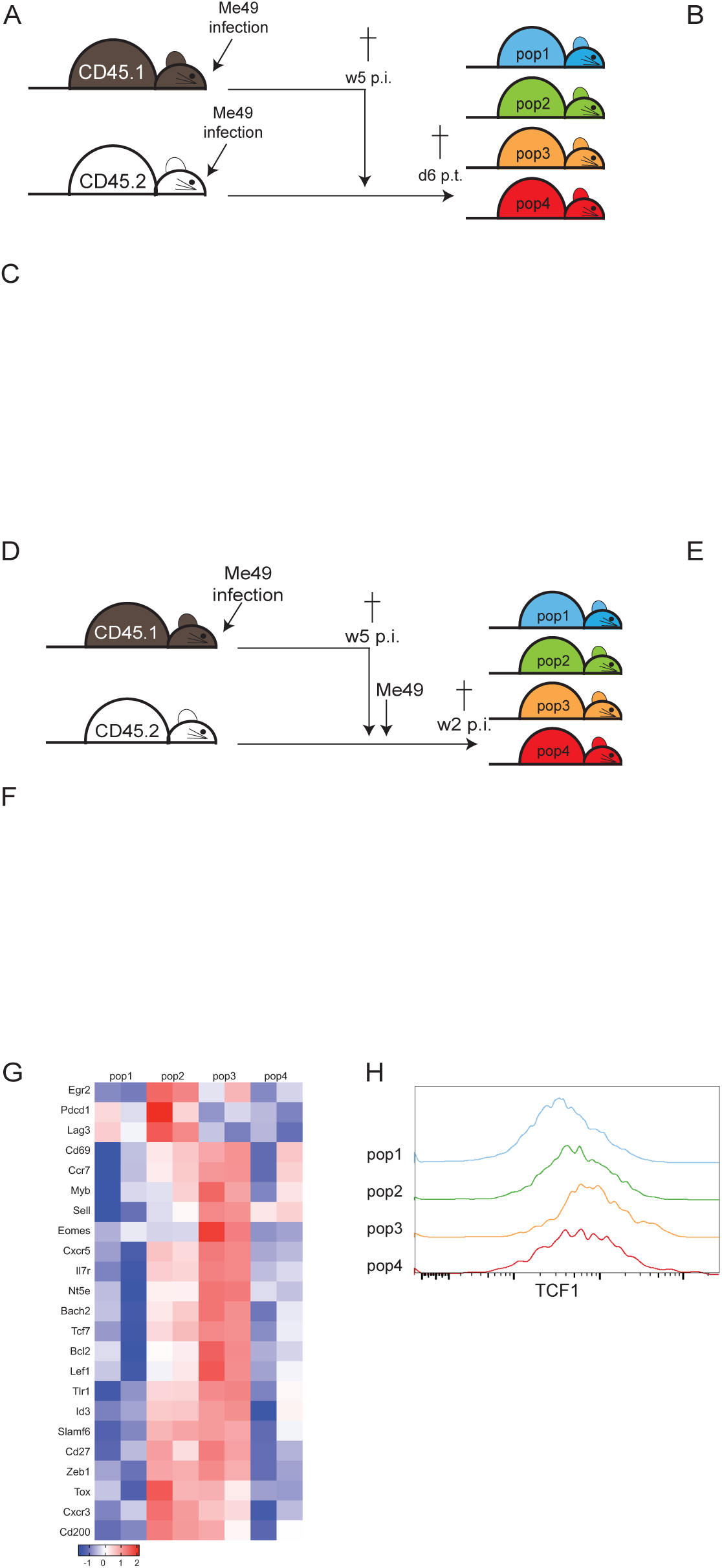
Stem-like memory CD8 T cell subset and its progeny follow an unconventional differentiation pathway when transferred to equally infected or naïve mice. Schematic for adoptive transfer of pop1-4 from 5 weeks infected CD45.1 mice to similarly infected CD45.2 animals. B) Number of CD45.1 cells recovered from the spleen of recipients at 6 days post-transfer. C) Phenotype of transferred cells recovered from each group of recipients (pop1-pop4 from left to right) presented as the frequency of CD45.1 cells. D) Schematic for adoptive transfer of pop1-4 from week 5 infected CD45.1 mice to naïve CD45.1 animals. Recipients were challenged one day post-transfer. E) Number of CD45.1 cells recovered from the spleen of recipients at day 15 p.i. F) Phenotype of transferred cells recovered from each group of recipients (pop1-pop4 from left to right) presented as the frequency of CD45.1 cells. G) Heatmap of DEGs associated with stem-like CD8 T cells in pop1-4 at week 5 p.i. H) gMFI of TCF-1 and Ly108 for pop1-pop4at week 5 p.i. Numbers on histograms represent the mean of gMFI for one experiment (4 mice) ± SEM for each marker. B, C, E, and F: Data presented as mean from individual mice ± SEM compiled from 2 experiments. *p<0.05, ***p≤0.001, and ****p<0.001 (one-way ANOVA with Bonferroni post-test).

Subsequently, the plasticity of CD8 T cell subsets in response to recall antigens was assessed by adoptively transferring the populations to naïve animals. The following day, the recipients were challenged with *T. gondii* cysts and the donor population was analyzed 14 days later (Fig. 3D). Similar to the previous experiment, CD45.1 cells from recipients transferred with pop1 retained their original phenotypes while an increased percentage of CD45.1 CD8 T cells from mice that received pop2 (compared to equally infected recipients) converted into pop1 (Fig. 3F). Akin to the transfer in equally infected mice (Fig. 3E), donor pop4 fully transitioned into pop1. Expectedly, CD45.1 cells recovered from mice transferred with pop3 (stem-like population) converted into every other phenotype. The ability of Pop3 to retain its original characteristics when transferred into a congenic infected recipient, as well as convert into pop1, pop2, and pop4 after challenge, further emphasizes its resemblance to stem-like CD8 T cells described in viral infection and cancer (33). Therefore, the transcriptome of pop3 (w5 p.i.) was compared to stem-like CD8 T cells from previously reported cancer and viral models (Table S2). As shown in figure 3 G, the similar expression pattern of multiple genes, including Tcf7, Bach2, Lef1, and Cxcr5, further highlights the close relationship of pop3 with the stem-like subset described in other systems (33). Additionally, flow cytometry analysis of TCF1 (encoded by Tcf7) and Ly108 (encoded by Slamf6), both typically increased in stem-like CD8 T cells (33), are also upregulated in pop3 (Fig. 3H). These findings demonstrate that, during chronic toxoplasma infection, pop3, with strong similarities to the stem-like CD8 T cells reported in chronic viral infections and cancer, is the only subset that can directly transition into other populations. Pop4, which resembles the intermediate effector memory CD8 T cells described in other models of exhaustion (34), is a transitory population derived from pop3 that converts exclusively into effector pop1.

### Memory CD8 T cell subsets display differential levels of inhibitory receptor expression

Studies from our laboratory have reported that similar to other systems, CD8 T cells from *T. gondii* infected mice show increased expression of inhibitory receptors like PD-1 during the chronic phase of the infection (15). In the present study, we observed that three memory (CD127^hi^) subsets (pop2-4) display higher expression of several inhibitory receptors (PD-1, TIGIT, LAG-3, 2B4, TIM-3, and CTLA-4) as compared to pop1 (Fig. 4A). Moreover, pop3 and pop4 display significantly higher expression of TIM-3, TIGIT, TIM3, 2B4, and CTLA-4 than pop2. Overall, the three memory subsets display elevated co-expression of multiple inhibitory receptors compared to effector pop1 (Fig. 4B). As previously reported in LCMV infection (35), an increase in PD-1 levels in pop2-4 was observed as early as two weeks p.i. (Fig. 4C), however, the decrease in PD-1, TIGIT, LAG-3, and 2B4 at the later time points could be caused by apoptosis of cells co-expressing multiple inhibitory receptors. Early expression of inhibitory molecules (PD-1, TIGIT, TIM3, LAG-3, CTLA4, and 2B4) was also observed in CD8 subsets from the brain and liver (Fig. S3A and B). However, expression of different inhibitory receptors by pop2-4 from the brain and liver remains constant throughout the course of the infection except for decreased expression of PD1 (brain and liver) and LAG-3 (liver only).

**Figure 4.**
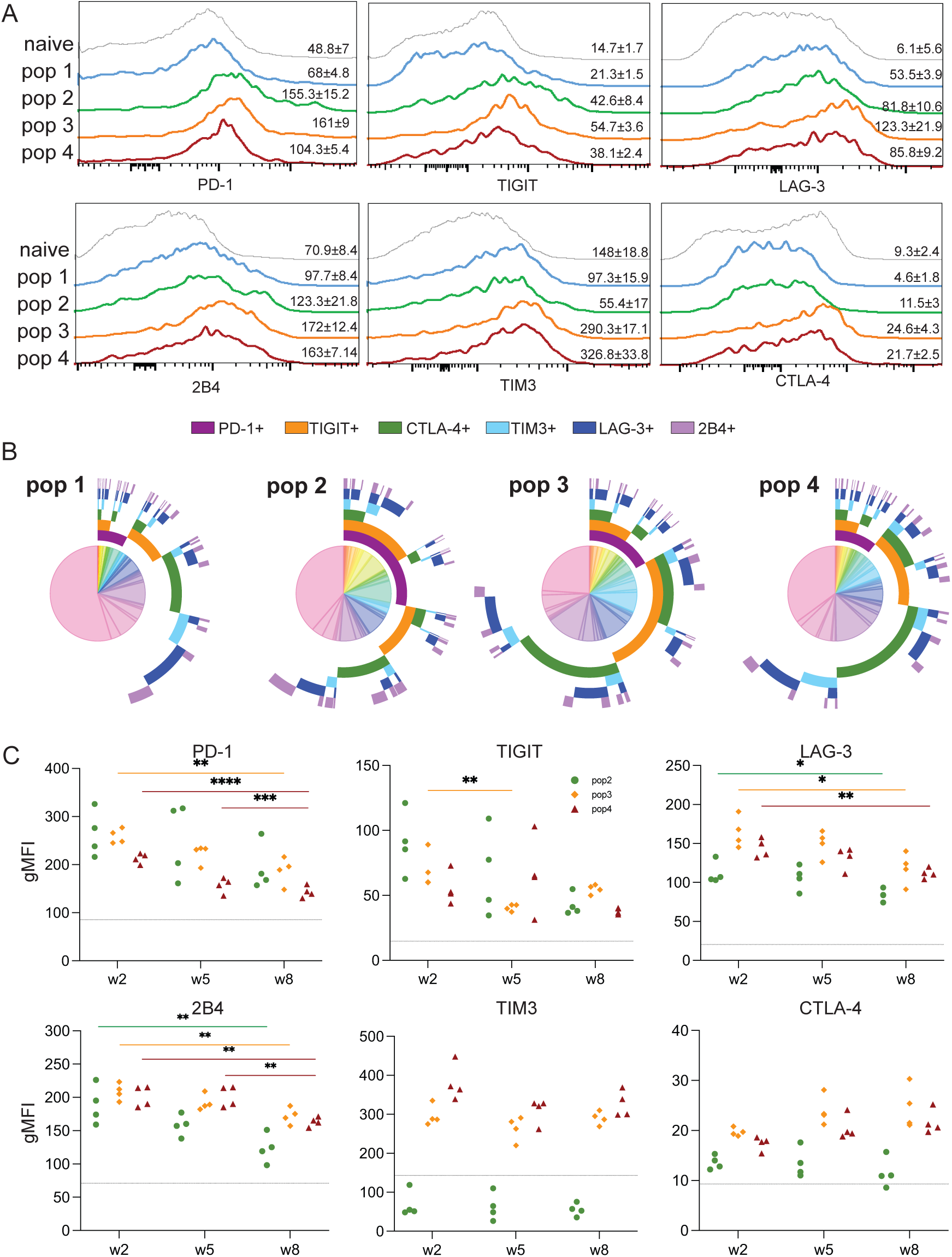
Increased multiple inhibitory receptor expression in stem-like memory CD8 subset (pop3) from chronically infected mice. Expression of PD1, TIGIT, LAG-3, 2B4, TIM3, and CTLA4 by pop1-pop4 from the spleen at week 8 p.i. Numbers on histograms represent the mean of gMFI for one experiment (4 mice) ± SEM of each marker in the four populations and CD8 T cells from naïve mice. B) Co-expression of multiple inhibitory receptors (PD1, TIGIT, LAG-3, 2B4, TIM3, and CTLA4) for pop1-pop4 at week 8 p.i. using SPICE. C) Kinetics of PD-1, TIGIT, LAG-3, 2B4, TIM3, and CTLA4 expression for pop2-pop4 at weeks 2, 5, and 8 p.i. gMFI of 4 individual mice is representative of 3 independent experiments. *p<0.05, **p<0.01, ***p≤0.001, and ****p<0.001 (one-way ANOVA with Bonferroni post-test).

CD8 T cell dysfunctionality during chronic viral infection is partially linked to the sustained presence of antigens (36). As replication of *T. gondii* takes place even during the chronic phase (27), the effect of continuous antigen persistence on the CD8 T cell subsets was assessed. To inhibit parasite replication, infected mice were treated with sulfamethoxazole (37) starting at week 6 p.i., and the CD8 T cell response was evaluated 2 weeks later. As shown in figure S4A, the frequency and numbers of Ag^+^ CD8 T cells are significantly lower when parasite replication is inhibited by antibiotic treatment, and the frequency of pop3 is increased while other populations are slightly reduced (Fig. S4B and C). However, inhibitory receptor expression by pop2-4 is not affected by the treatment except for a significant decrease in TIGIT expression by pop2 (Fig. S4D).

As previously described, a high antigen load early after infection promotes the differentiation of precursor T cells with acquired hallmarks of exhaustion (38). Moreover, a recent study from our laboratory reported that CD8 T cell memory generation against *T. gondii* infection is dose-dependent, with a lower dose driving a stronger Ag^+^ CD8 T cell response at week 8 p.i. (39). In the current study, pop1, pop2, and pop4 cell numbers are increased in mice infected with 2 cysts compared to animals receiving a 10-cyst challenge (Fig. S5A and B). Higher numbers in pop2 from 2 cysts infected mice could be explained by lower PD1 expression (Fig. S5C). However, the increase in pop1 and pop4 in these animals may be caused by enhanced pop3 conversion. Thus, although a lower dose of infection generates a superior CD8 T cell response, it is not correlated with a decreased expression of inhibitory receptors.

Our data demonstrate that stem-like pop3, intermediate effector memory pop4, and pop2 exhibit elevated inhibitory receptor expression starting as early as week 2 p.i. However, this increase in multiple receptor expression is not dependent on the dose of antigen during priming or the chronic phase of infection. Moreover, during the chronic phase of the infection, pop3 is more predominant when the antigen load is reduced by antibiotic treatment since effector pop1 is no longer needed, making pop3 conversion redundant.

## Intermediate effector/memory population exhibits enhanced functionality

CD8 T cells, an important source of IFNγ during *T. gondii* infection, are crucial for controlling the dissemination of the parasites and for protection during acute and chronic infection (13). Additionally, cytolytic activity against infected targets is critical, especially during the chronic stages of the infection (15, 40). As expected, at week 2 p.i., all subsets are in cell cycle (based on proliferation marker Ki67) (Fig. 5A), and stem-like pop3 exhibits moderate proliferation compared to the other subsets. While the reduction in expansion of pop1, pop2, and pop4 during the chronic phase is anticipated, stem-like pop3 proliferation decreases rather unexpectedly (Fig. 5B and C). Importantly, a fraction of pop4 and stem-like pop3 remains in cell cycle, as demonstrated by an increase in Ki67 MFI in the Ki67^+^ cells at week 8 p.i. (pop3: 2611±69 (w2), 4965±1051(w8) and pop4: 3485±241 (w2), 6220±2000 (w8) ANOVA with Bonferroni post-test: p<0.01). In accordance with our RNA sequencing data (Fig. 2F), pop1 and pop4 exhibit significantly higher IFNγ production and cytotoxic activity (CD107^+^) at week 2 p.i., as compared to pop3 (Fig. 5E and F). Interestingly, while there is a reduction in the percentage of bi-functional CD8 T cells within pop1 and pop2 at week 8 p.i., pop4 remains cytotoxic and can still produce IFNγ and TNFα (Fig. 5F). The sustained functionality of pop4 during chronic toxoplasmosis could be attributed to the continued upregulation of several costimulatory molecules (Fig. S6A-B), that are known to play an important role in T cell activation (41). As shown in figure S6C, pop4 continues to exhibit increased expression of multiple costimulatory molecules even during the later stages of chronic infection.

**Figure 5.**
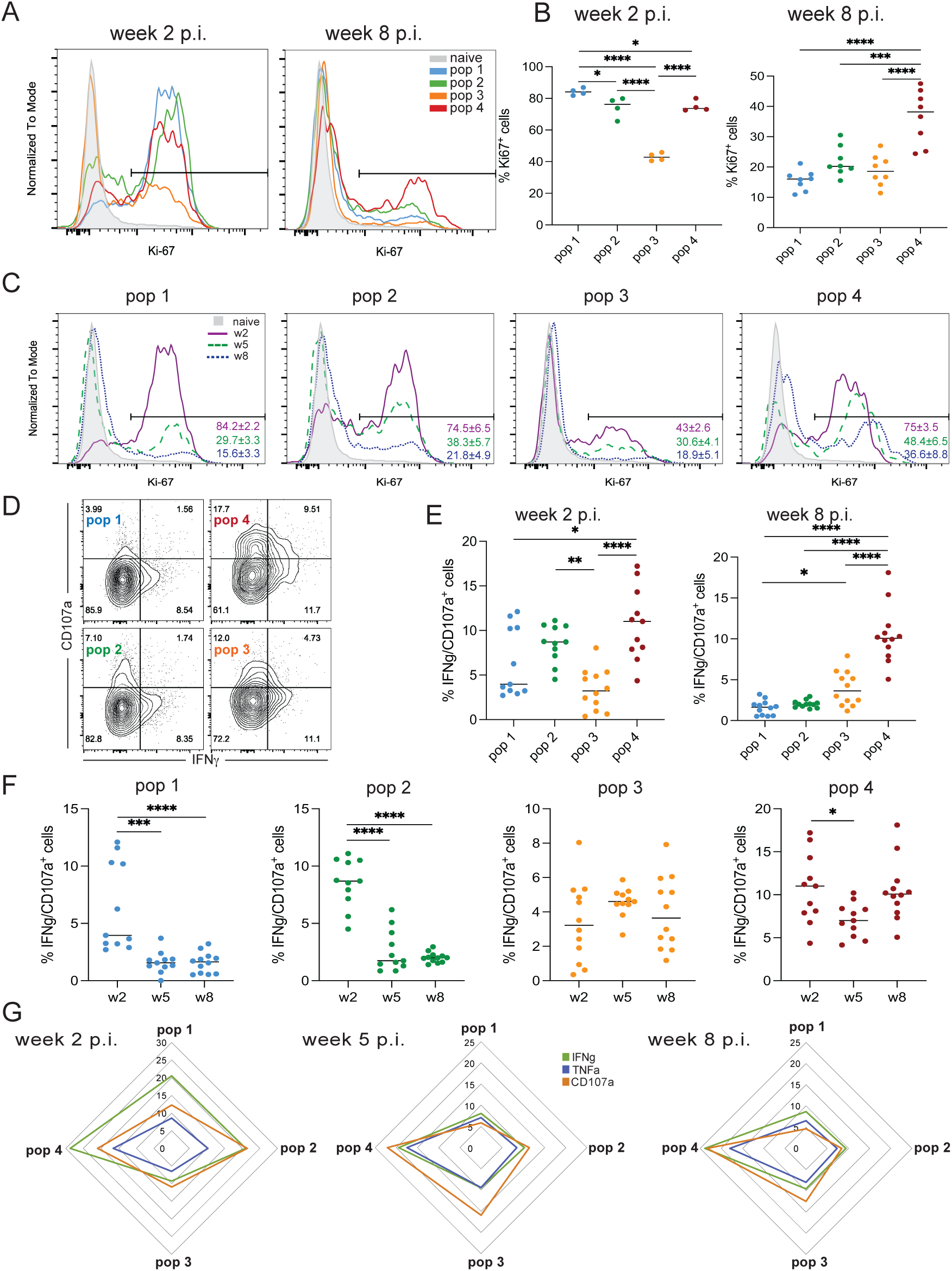
Intermediate effector/memory pop4 maintains elevated functional fitness during chronic *T. gondii* infection. A) Ki67 expression for pop1-pop4 from the spleen at weeks 2 p.i. and 8 p.i. CD8 T cells from naïve mice are used as a control. B) Frequency of Ki67^+^ cells in pop1-pop4 at week 2 and 8 p.i. C) Kinetic of Ki67 expression in pop1-4 at weeks 2, 5, and 8 p.i. Numbers in the graph represent the mean percentage of Ki67+ cells for one experiment (4 mice) ± SEM. D) Intracellular staining for IFNγ and CD107a in pop1-pop4 at week 5 p.i. E) Percentage of IFNγ^+^CD107a^+^ cells from pop1-pop4 at weeks 2 and 8 p.i. F) Kinetics of the frequency of IFNγ^+^CD107a^+^ cells from pop1-pop4 at weeks 2, 5, and 8 p.i. G) Spider plot for pop1-pop4 shows the frequency of IFNγ^+^, TNFα^+^, and CD107a^+^ cells from the different populations at weeks 2, 5, and 8 p.i. B, E, and F: Data presented as the mean for individual mice ± SEM compiled from at least two experiments. *p<0.05, **p<0.01, ***p≤0.001, and ****p<0.001 (one-way ANOVA with Bonferroni post-test).

In conclusion, while the decrease in the proliferative ability of pop1, pop2, and pop4 is not surprising, the unexpected reduction in Ki67^+^ pop3 could be correlated to the elevated co-expression of multiple inhibitory receptors. As seen in other models, the intermediate pop4 displays strong functionality throughout the course of the infection that could be explored for immunotherapeutic purposes.

## Discussion

Polyfunctional CD8 T cells are critical for controlling *T. gondii*, especially during the chronic phase of the infection (13). However, in an encephalitis model, reactivation of chronic infection attributed to the dysfunctional state of memory CD8 T cells leads to severe encephalitis and host mortality (15). Cues from chronic viral infections and tumor models, where CD8 T cells play a pivotal role in protection, indicate that memory CD8 T cells are a heterogeneous population with different receptor expression, functional potential, migration, and tissue localization (4). As previously reported (23) and based on our data, *T. gondii* CD8 T cell response does not conform to the T_CM_/T_EM_ model. Similarly, our populations could not be defined based on CD27 and a glycoform of CD43 as reported in a respiratory viral infection (42). While we used a clustering algorithm to define our populations, Wilson *et al.* had previously used the same markers to study the role of IL-12 in the CTL response against *T. gondii* infection (22). This report demonstrated that CTL cells can be fractionated into a T_CM_ fraction I (KLRG1^-^ CD62L^hi^), T_EM_ fraction II (KLRG1^-^CD62L^lo^), effector cell rich fraction III (KLRG1^+^CD62L^lo^), and fraction IV (KLRG1^+^CD62L^hi^) during both acute and chronic phases of infection. However, these fractions were not characterized beyond KLRG1 and CD62L expression. In our study, pop1 displays the characteristics of terminally differentiated effector CD8 T cells (KLRG1^+^CD62L^lo^) with significant increases in molecules associated with cytotoxicity, cytokines, and various annexin and caspases (Fig. 2F). Pop1 remains the most dominant subset even though its frequency decreases as the infection progresses to chronicity. Pop2 (KLRG1^-^CD62L^lo^) is similar to T_EM_ with low levels of KLRG1, CD62L, CCR7, and high inhibitory receptor expression and pop3 (KLRG1^-^CD62L^hi^) expresses multiple genes associated with stem-like memory CD8 T cells. Pop4 (KLRG1^+^CD62L^hi^) displays an intermediate effector/memory profile with elevated functionality and strong expression of costimulatory molecules even during the chronic phase of the infection. Interestingly, pop4 also exhibits high inhibitory receptor expression that could be inherited from pop3, as adoptive transfer studies demonstrate that this population is directly derived from the stem-like memory subset. These adoptive studies also highlight the transient nature of pop4, as this population converts into effector pop1 instead of a terminally exhausted subset as described in chronic viral and cancer models (4, 34). Even though the transient pop4 phenotypically resembles pop3, RNA seq analysis demonstrates that this subset is more similar to pop1 than parental pop3.

Similar to the stem-like CD8 T cells in viral and cancer models, pop3 demonstrates increased expression of Tox, Tcf1, Bach2, Lef1, Id3, Slamf6, and Zeb1, as well as genes related to translation, especially those encoding ribosomal proteins (32). More importantly, adoptive transfer studies demonstrate that pop3 retains its phenotype when transferred to equally infected mice but converts into pop2, pop4, and possibly pop1 when transferred to naïve mice that were subsequently challenged. However, in chronic viral and cancer models of CD8 T cell exhaustion, this self-renewing PD-1^+^ TCF-1^+^ population contributes to the maintenance of robust T cell immunity under chronic antigenic stimulation (11, 43). Due to their long-term persistence and ability to transition into different subsets, they are considered ideal targets for cancer immunotherapy. As PD-1 blockade accelerates the differentiation of stem-like memory CD8 T cells into an effector population (44), pop3 could play a critical role in controlling the reactivation of chronic *T. gondii* infection since CD8 T cells can be rescued by anti-PDL-1 treatment (15).

Interestingly, pop4, derived from pop3, retains some levels of commonality with the parental stem-like memory subset based on the shared expression of several surface markers like CD62L, CD127, CCR7, and CD27. When adoptively transferred, the majority of pop4 transitioned into pop1, and these two populations are very similar, differing only by 43 DEGs with Sell (which encodes CD62L) being the most dysregulated gene. Notably, pop4 displays some of the attributes of both effector (KLRG1^+^) and memory (CD127^hi^) CD8 T cells akin to the hybrid effector-memory phenotype described by Chu *et al* in a resistant model of chronic toxoplasmosis (23). In this study, CXCR3^-^KLRG1^-^ CD8 T cells were identified as effector CD8 T cells, CXCR3^+^KLRG1^-^ as a memory subset, and CXCR3^+^KLRG1^+^ as an intermediate effector/memory population with antigen-dependent maintenance. In our model, based on CXCR3 and KLRG1 expression by the different populations, pop2, and pop3 resemble the CXCR3^+^KLRG1^-^ subset, while pop4 bears a striking likeness to the intermediate CXCR3^+^KLRG1^+^ population (23). However, unlike our findings, none of the subsets described in this report expressed CD62L or CCR7. In our study, the elevated functionality of intermediate effector memory pop4, which has been reported in other models (11, 34), is correlated with high inhibitory receptors and costimulatory molecule expression. Interestingly, pop4 exhibits a significantly higher frequency of IFNγ^+^CD107a^+^ than pop1 as early as week 2 p.i. Our data, combined with the earlier report from Chu *et al* (23) in a resistant model of *T. gondii* infection indicates that this intermediate effector/memory population could control parasite reactivation. However, in susceptible C57BL/6 mice, even though highly functional pop4 is continuously generated, their number is eclipsed by less functional pop1. This imbalance continues even during the chronic phase of the infection, potentially impairing their ability to control parasite reactivation in a susceptible model.

Overall, the CD8 T cell memory response during chronic toxoplasmosis in an encephalitis model appears to be a complex process with different subsets exhibiting various states of exhaustion and functionality. To date, most of the studies on CD8 T cell exhaustion have focused on cancer and chronic viral infection models. A consensus has emerged in these systems, and the different subsets of CD8 T cells are defined as stem-like exhausted progenitor CD8 T cells (Tex-prog), exhausted intermediate phenotype (Tex-int), and highly exhausted terminally differentiated effector (Tex-eff) (11, 45). Tex-prog converts into Tex-int, which then differentiates into Tex-eff. Based on our studies, memory CD8 T cell subsets during chronic toxoplasmosis exhibit a distinctive pattern of differentiation. In our model, pop3 converts directly into pop4, pop2, and potentially pop1 instead of following the linear transition pattern from Tex-prog into Tex-int and finally Tex-term described in cancer and chronic viral systems (45). Even though these memory CD8 T cell populations retain some commonalities across different infections, differences can be driven by pathogen-specific cues. Evidence that the parasites continue to replicate (27) and terminally differentiated pop1 still encompasses the majority of the CD8 T cells during chronic toxoplasmosis should be taken into consideration. Other complex parasitic infections like malaria and Leishmania infection, where CD8 T cell exhaustion has been reported (18, 46), could also provide vital clues into the lineage of memory CD8 T cell differentiation. Similarly, CD8 T cells with stem-like properties have been reported in nonviral/cancerous models, but their transition during the state of exhaustion has not been determined (18, 19). The discrepancy between the subsets described in our study and other well-described systems of CD8 T cell exhaustion demonstrates that this phenomenon is more complex, and the mechanisms driving CD8 T cell fate decisions during different chronic infections need to be better characterized.

## Materials and methods

### Mice and parasites

C57BL6/J and B6-Ly5.2 (CD45.1) mice were purchased from The Jackson Laboratory and housed in a specific pathogen-free facility at The George Washington University. All experiments were performed per institutional policies and regulations of the relevant animal welfare acts and have been reviewed and approved by The George Washington University institutional animal care and use committee. *T. gondii* cysts of the Me49 strain were isolated from the brains of infected CBA/J mice at 5 weeks post-infection. Six to eight-week-old mice were infected via the intragastric route with 2 or 10 cysts. Mice were sacrificed by CO2 inhalation at different time points post-infection. *T. gondii* lysate antigen (TLA) was prepared from the RH strain of *T. gondii* as previously described (21). For some experiments, Toxoplasma-infected mice (6 weeks p.i.) were administered sulfamethoxazole (800 mg/l) in drinking water for 2 weeks (37), and medicated water was replaced every third day.

### Lymphocyte isolation and staining

Single-cell suspension was prepared from the spleen, liver, blood, and brain of mice using standard protocols (21). Briefly, splenic single-cell suspension was made by mechanical disruption followed by red blood cell lysis. The liver was perfused and dissociated with a mesh sieve before hepatic lymphocytes were enriched using a 30% Percoll gradient. The brain was harvested and passed through a 70µm cell strainer, and the single-cell suspension was centrifuged on a 30% Percoll. Peripheral blood was collected in PBS with 100U/ml heparin, and partial red blood cell lysis was carried out. Lymphocytes were stained with fixable live/dead dye followed by surface staining for 20 minutes at 4^0^C in PBS supplemented with 2% bovine calf serum with the following antibodies: CD8 (YTS156.7.7), CD11a (M17/4), CD44 (IM7), CD62L (MEL-14), KLRG1 (2F1), CD43 (1B11), CD27 (LG.3A10), CCR7 (4B12), CX3CR1 (SA011F11), CXCR5 (L128D7), CXCR3 (CXCR3-173), CTLA-4 (UC10-4B9), TIGIT (1G9), 2B4 (m2B4(B6)458.1, TIM3 (RMT3-23), LAG-3 (C9B7W), PD-1 (RMP1-30), CD28 (37.51), OX40 (OX-86), CD40L (MR1), GITR (DTA-1), CD45.1 (A20), and CD45.2 (104) were purchased from Biolegend and CD127 (A7R34), CD27 (LG.7F9), 41BB (17B5), ICOS (7E.17G9), and Ly108 (13G3-19D) from Thermofisher Scientific. Tgd057 tetramers were obtained for the NIH tetramer facility (Atlanta, GA). Tetramer staining was performed at 4^0^C for 1 hour in PBS supplemented with bovine calf serum.

For intracellular cytokine detection, lymphocytes were stimulated for 18 hours with 30µg/ml of TLA, and Protein Transport Inhibitor Cocktail (eBioscience, Thermofisher Scientific) was added during the last 4 hours of restimulation. After surface staining and permeabilization/fixation (IC Fixation buffer, Thermofisher Scientific), cells were stained for intracellular cytokine detection (anti-IFNγ (XMG1.2) from Biolegend and anti-TNFα (MP6-XT22) from Thermofisher Scientific) as previously described (21).

Staining for transcription factor detection (anti-Ki67 (SolA15) from Thermofisher Scientific and anti-TCF1 (C63D9) from Cell Signaling) was performed after surface staining and a permeabilization/fixation step using Cytofix/Cytoperm kit (BD Biosciences).

Data were acquired with a BD FACSCelesta or LSR Fortessa and analyzed with FlowJo software (BD Biosciences). Dead cells were systematically excluded using the LIVE/DEAD fixable aqua dead cell stain kit (ThermoFisher Scientific), and quality control was performed with FlowAI (FlowJo). Co-expression analysis was performed with SPICE 6 (47). Multiparametric flow cytometry data were analyzed and visualized with FlowSOM and TriMap dimensionality reduction algorithms.

### Adoptive transfer

CD8 T cells were purified from 5-7 pooled spleens of CD45.1 infected mice using the EasySep Mouse CD8 T cell isolation kit (Stemcell Technologies). The enriched CD8 cell suspension was then labeled with fluorescent antibodies against CD8, CD11a, CD44, CD62L, KLRG1, and the different populations were then sorted with a BD Influx cell sorter (purity ≥90%). Dead cells were excluded using Sytox Blue (ThermoFisher Scientific). Cells from each population were individually transferred to infected or naïve CD45.2 mice (10^5^ cells/mouse) via tail vein injection. Naïve recipient mice were challenged the next day with *T. gondii* (10 cysts/mouse) via intragastric route.

### RNA extraction, sequencing, and analysis

CD8 T cell populations pooled from 5-7 mice (week 5 and 8 p.i.) were sorted as described above. RNA extraction was performed using the RNeasy Plus mini kit (Qiagen) following the manufacturer’s protocol. Each sample group was comprised of two experimental replicates. Ribosomal RNA was removed using a NEBNext rRNA Depletion kit, followed by library preparation with NEBNext Ultra II RNA Library Preparation kit for Illumina (New England BioLabs). Samples were sequenced by Genewiz (Azenta Life Sciences) on an Illumina HiSeq platform with 2x150 bp sequencing read length. RNAseq data are available in the GEO database (accession GSE280150). RNA-seq analysis was performed using Galaxy (48). After quality control and adapter sequences trimming (Cutadapt Galaxy version 4.7+galaxy0) (49), the remaining rRNA was removed using SortMeRNA (Galaxy version 2.1b.6)(50). Reads were mapped to the mouse genome (GRCm38/mm10) using RNA STAR (Galaxy version 2.7.11a+galaxy0) (51). GC bias was evaluated with computeGCBias (Galaxy version 3.5.4+galaxy0) and corrected when needed using correctGCBias (Galaxy version 3.5.4+galaxy0) (52). Transcripts were quantified using featureCounts (Galaxy version 2.0.3+galaxy2)(53), and differential gene expression analysis was performed using DESeq2 (Galaxy version 2.11.40.8+galaxy0)(54). Genes were considered differentially expressed if they achieved an adjusted p-value <0.05 and |log2 fold change| >1. PCA was generated from VST-normalized counts using PCA plot w ggplot2 (Galaxy version 3.4.0+galaxy0) (55). The volcano plot and Venn diagram were generated in Galaxy (Galaxy version 0.0.5 and Galaxy version 0.1.1, respectively). Spearman’s Correlation Coefficient plot was generated with log-transformed normalized count data generated by DESeq2. The top 3800+ DEGs among the four populations (pop1-4) were selected for further analysis and divided into 6 clusters by K-means clustering (numeric clustering, Galaxy version 1.0.11.0). The data was visualized with heatmaps using the z-scores and log2(+1)-transformed normalized counts. Overrepresented gene ontology categories were determined with Metascape (https://metascape.org)(56). Spearman’s correlation and heatmap were visualized using GraphPad Prism (Version 10.2.2).

### Statistical analysis

Differences in percentage, absolute number, and MFI for each experiment were assessed using one-way ANOVA with Bonferroni post-test. Error bars in the graph indicate the SEM of values from individual mice. All computations were completed using GraphPad Prism software.

## Acknowledgments

We thank Gregory Cresswell and the Flow Cytometry Core Facility at George Washington University for assistance with sorting of the CD8 T cell subsets. The core is supported by the George Washington University Cancer Center and School of Medicine and Health Sciences.

## Funding

This work was supported by the National Institute of Health grants AI15308 and AI33325 awarded to IK.

## Notes

### Competing Interest Statement

The authors have declared no competing interest.

### Summary of Updates

Manuscript has been revised for improved clarity. Title and abstract have been modified to emphasize the novelty of the findings

