## Supplementary figures and images for "In a model of parasite-mediated exhaustion, stem-like CD8 T cells differentiate into an unconventional intermediate effector memory subset"

### supplementary figure 1

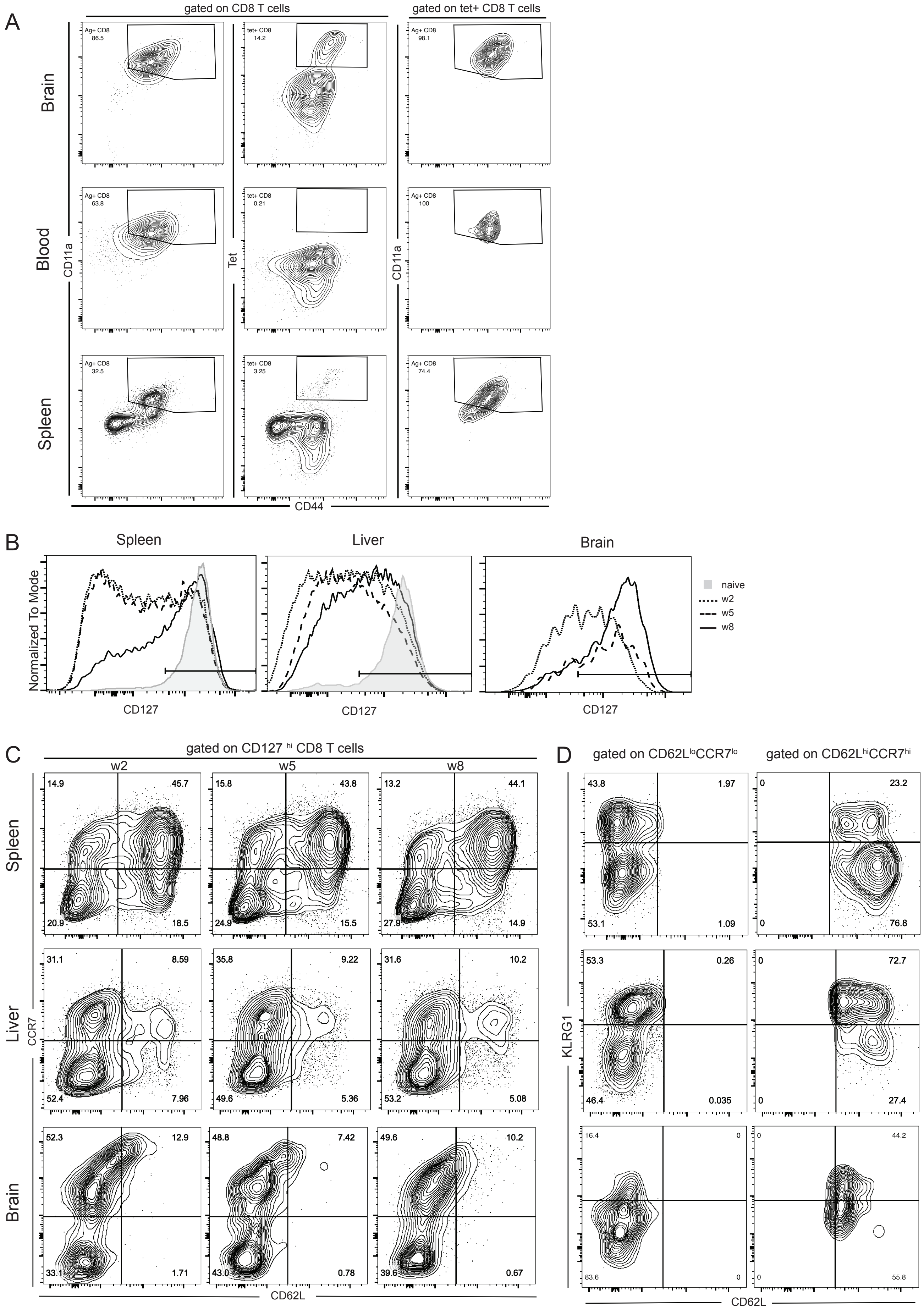

### supplementary figure 2

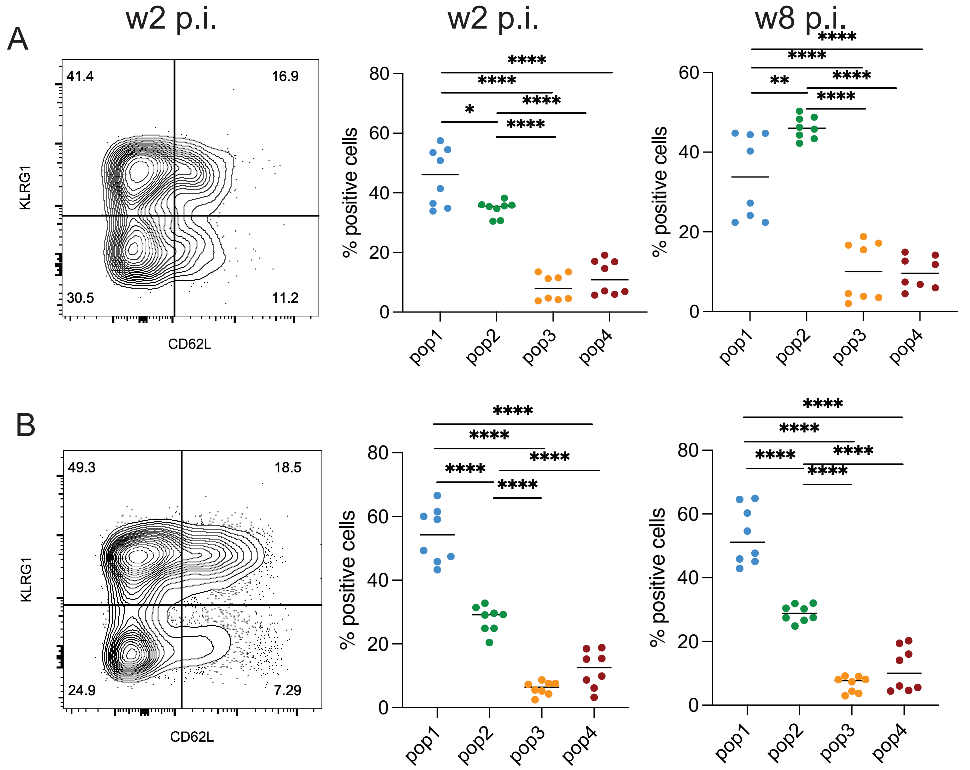

### supplementary figure 3

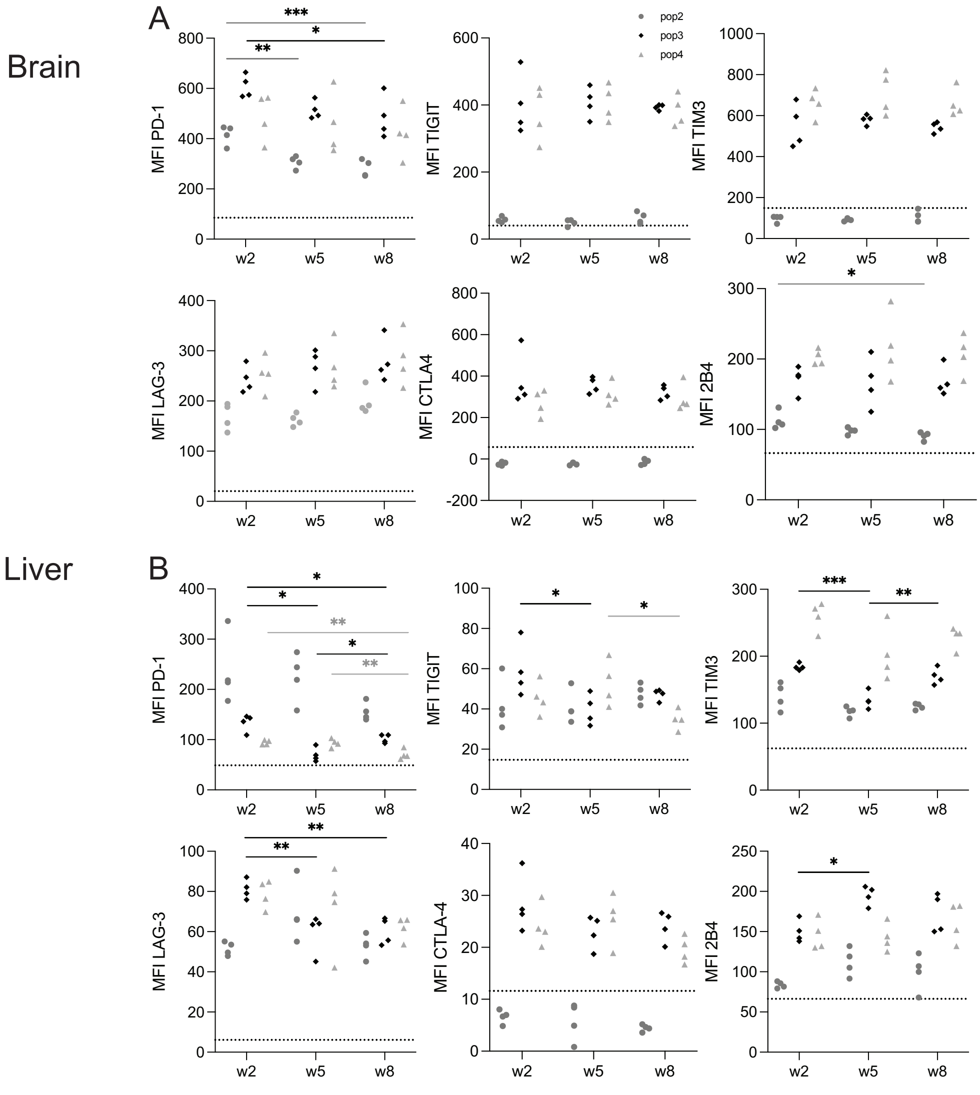

### supplementary figure 4

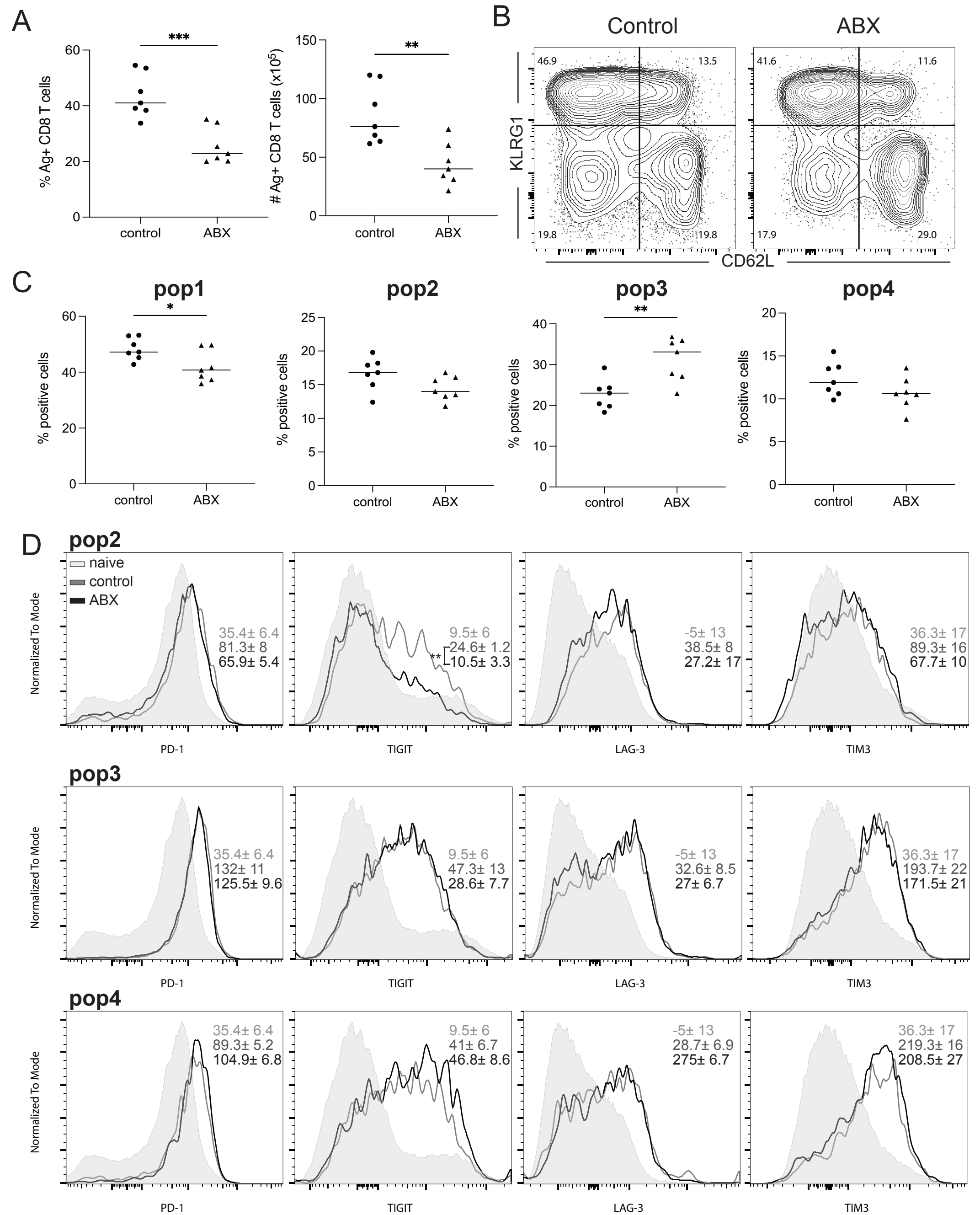

### supplementary figure 5

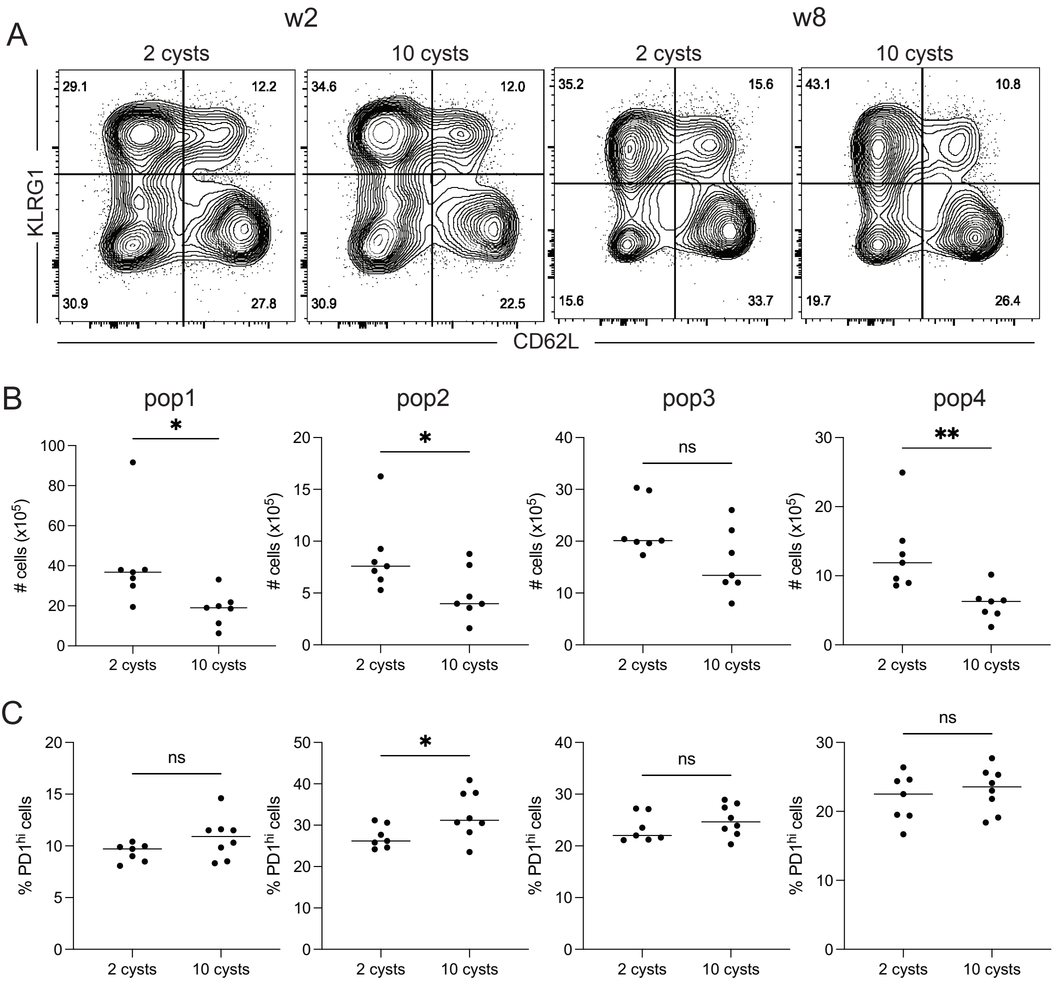

### supplementary figure 6

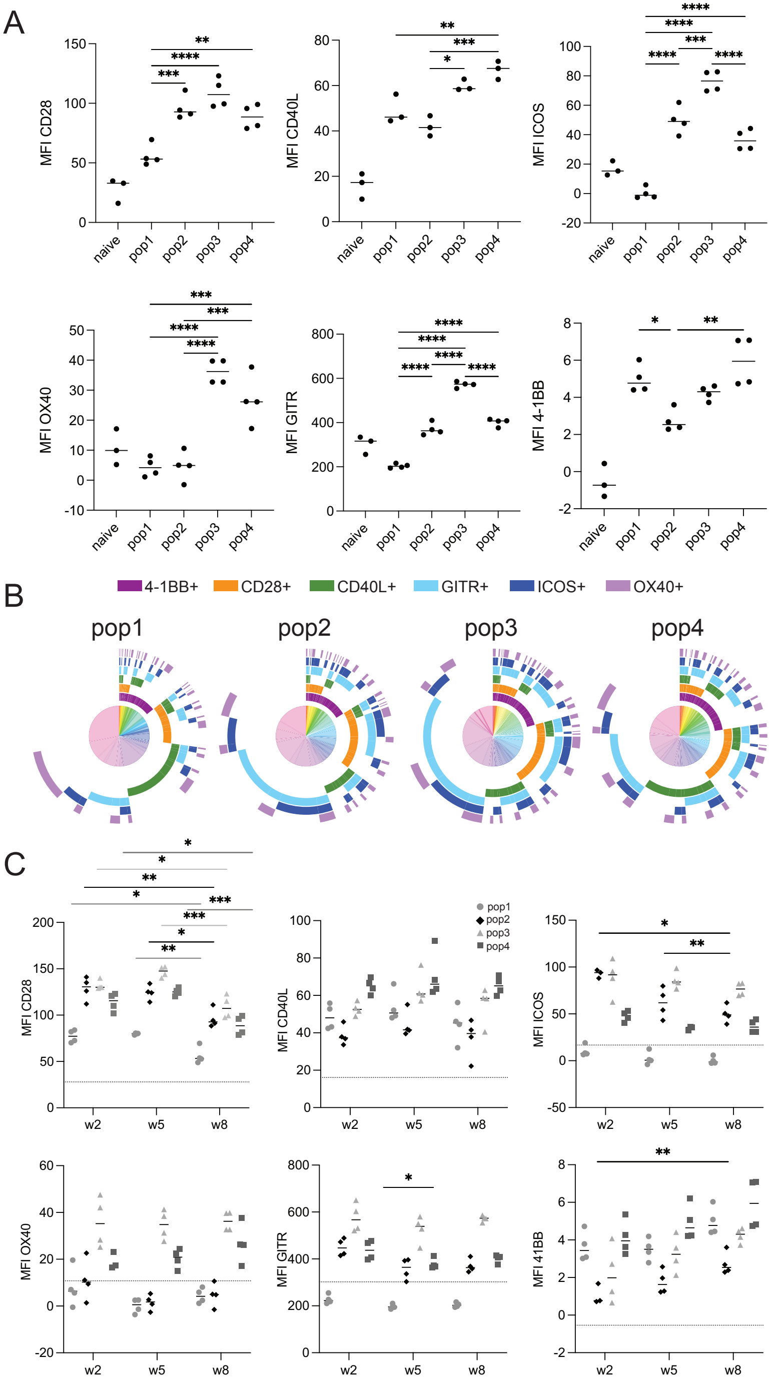

### supplementary table 2

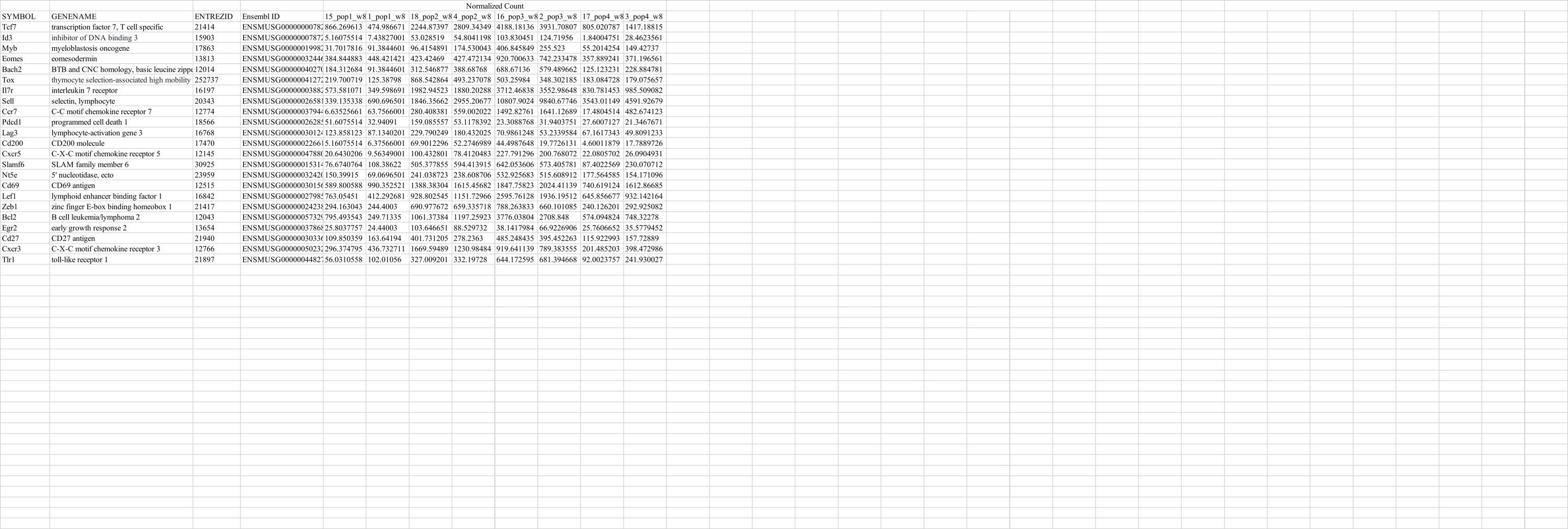
